# Concrete waste as a source of calcium and alkalinity for coccolithophore cultivation: an experimental test of carbon dioxide removal and sequestration

**DOI:** 10.64898/2026.10.31.763296

**Authors:** Maxime Ferriere, Antonin Affholder, Jack Welchert, Matthew Katterman, Hang Zeng, Hee-Jeong Kim, Daniel Apai, Joel Cuello

## Abstract

Cultivation of calcifying microalgae has been proposed as a route to carbon dioxide removal (CDR), but biogenic calcification releases CO_2_ unless the calcium supplied also adds alkalinity. We used cultures of the coccolithophore *Chrysotila* sp to test whether hydrated lime (Ca(OH)_2_), supplied as a reagent or within ground waste concrete, can provide both calcium and alkalinity. In a 26-day factorial batch experiment, we monitored pH, nitrate and dissolved calcium. Calcite was quantified by thermogravimetric analysis. Nitrate concentration decreased and calcite accumulated in all algal treatments, indicating that lime or concrete did not inhibit growth or calcification. Lime and concrete raised culture pH significantly whereas the algal growth rate inferred from nitrate drawdown did not differ among treatments. Calcite production was 3.6-fold higher with lime than in seawater (4.5 vs. 1.2 mg L^−1^ d^−1^) and highly variable with concrete; lime and concrete cultures remained supersaturated with respect to calcite (Ω ≈ 7-10 vs. 0.2). Carbonate-system calculations indicate that the added alkalinity raises the equilibrium CO_2_ uptake capacity of the medium by *ca.* 3.4 mmol kg^−1^, offsetting CO_2_ released by calcification. These results provide proof of concept for coupling coccolithophore cultivation with concrete waste reuse for CDR.

## 1 INTRODUCTION

Limiting global warming to well below 2 °C will require both deep reductions in greenhouse gas emissions and the large-scale removal and durable storage of atmospheric carbon dioxide (US EPA, 2025; Riahi et al., 2022). Engineered approaches to CO_2_ removal (CDR), such as direct air capture, remain expensive and difficult to scale (Rissman et al., 2020), motivating the evaluation of nature-based alternatives. Because the ocean accounts for about 70% of the Earth’s surface–atmosphere interface, ocean-based processes are likely to be part of any cost-effective and scalable CDR portfolio (NASEM, 2022).

Coccolithophores are unicellular calcifying marine algae that fix dissolved inorganic carbon into organic biomass by photosynthesis and into calcium carbonate (calcite, CaCO_3_) by intracellular calcification, producing calcite scales called coccoliths (Billard & Inouye, 2004). They are globally widespread and form blooms that are visible from space (Sadeghi et al., 2012). Over geological time, coccolithophores have had a large impact on the carbon cycle: the chalk cliffs of Dover (UK) and Normandy (France) formed largely from coccoliths deposited during the Late Cretaceous, and coccolithophores remain key players in the present-day marine carbon cycle (de Vargas et al., 2007). The fertilization of natural blooms and the mass cultivation of calcifying microalgae in photobioreactors have therefore been proposed as strategies for carbon capture and sequestration (Jansson & Northen, 2010; Apai et al. 2024).

A counter-intuitive obstacle stands in the way. Calcifying algae do not take up CO_2_ from the atmosphere directly; they use bicarbonate (HCO_3_^−^) and carbonate (CO_3_^2−^) ions already present in seawater. The precipitation of calcite consumes two moles of alkalinity per mole of CaCO_3_ formed and returns CO_2_ to the water and, ultimately, to the atmosphere (Frankignoulle et al., 1994; Bialik et al., 2022):

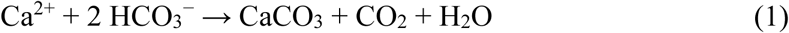

Thus, if the calcium used by the algae is supplied without alkalinity, calcification acts as a net source of CO_2_, the opposite of the intended outcome. Net CDR requires that calcium be supplied together with alkalinity, so that the ocean–atmosphere solubility pump draws additional CO_2_ into the water (Table 1). Most calcium used industrially is mined as gypsum (CaSO_4_), whose dissolution adds no alkalinity. In contrast, hydrated lime (portlandite, Ca(OH)_2_) dissociates into Ca^2+^ and OH^−^ and raises alkalinity by two equivalents per mole of calcium. Hydrated lime is commercially available and is also present in substantial amounts in concrete. Concrete waste, generated in large quantities by demolition, might therefore provide an inexpensive and sustainable source of both calcium and alkalinity for coccolithophore cultivation.

**Table 1.** Net effect on CO_2_ in the atmosphere, the water and the mineral phase of the main processes involving one mole of calcium.

| **Process** | **Reaction** | **Atmosphere** | **Water (DIC)** | **Minerals** |
|---|---|---|---|---|
| Calcite precipitation | Ca<sup>2+</sup> + 2 HCO<sub>3</sub><sup>−</sup> → CaCO<sub>3</sub> + CO<sub>2</sub> + H<sub>2</sub>O | +1 | −2 | +1 |
| Alkalinization by hydrated lime | Ca(OH)<sub>2</sub> + 2 CO<sub>2</sub> → Ca<sup>2+</sup> + 2 HCO<sub>3</sub><sup>−</sup> | −2 | +2 | 0 |
| Alkalinization + biocalcification | Ca(OH)<sub>2</sub> + CO<sub>2</sub> → CaCO<sub>3</sub> + H<sub>2</sub>O | −1 | 0 | +1 |
| Calcination (cement production) | CaCO<sub>3</sub> → CaO + CO<sub>2</sub> | +1 | 0 | −1 |
| Calcite dissolution (weathering) | CaCO<sub>3</sub> + CO<sub>2</sub> + H<sub>2</sub>O → Ca<sup>2+</sup> + 2 HCO<sub>3</sub><sup>−</sup> | −1 | +2 | −1 |

*Note*: Values are moles of CO<sub>2</sub> added (+) or removed (−) from each compartment per mole of Ca<sup>2+</sup> involved; carbon in the water is counted as dissolved inorganic carbon.

Table 1 summarizes the net effect on CO_2_ in the atmosphere, the water and the mineral phase of the main processes involving one mole of Ca^2+^. Alkalinization alone (adding Ca(OH)_2_ without calcification) removes two moles of CO_2_ from the atmosphere but stores it as dissolved carbonate species, whose permanence depends on the long-term fate of the water. Alkalinization followed by biocalcification removes only one mole of atmospheric CO_2_, but stores it in calcite, a stable mineral that could also serve as a biogenic substitute for limestone. This trade-off makes it important to determine whether coccolithophores can grow and calcify efficiently when calcium and alkalinity are supplied as hydrated lime, and whether concrete waste can provide hydrated lime in a usable form.

Here we report a first experimental test of this idea using the coccolithophore *Chrysotila* sp. We hypothesized that ground concrete waste, which contains calcium hydroxide, increases CO_2_ solubility by raising alkalinity and provides a sustainable source of calcium for growing coccolithophores, enabling net removal and sequestration of atmospheric CO_2_. We tested four predictions: supplementing calcium as hydrated lime (P1) does not inhibit algal growth or calcium uptake; (P2) increases pH, which favors algal growth; (P3) increases alkalinity and therefore atmospheric CO_2_ solubility; and (P4) increases calcite production and the calcite saturation state, maintaining conditions favorable to calcification. For each prediction, we asked whether the effect persists when hydrated lime is supplied within concrete waste.

## 2 MATERIALS AND METHODS

### 2.1 Organism and culture medium

A strain of the coccolithophore genus *Chrysotila* (strain AC237) was obtained in 2023 from the Roscoff Culture Collection (CNRS, Station Biologique de Roscoff, France) and maintained in the laboratory on the standard culture medium. The medium consisted of artificial seawater (ASW; 450 mmol L^−1^ NaCl, 10 mmol L^−1^ KCl, 9 mmol L^−1^ CaCl_2_, 30 mmol L^−1^ MgCl_2_ and 16 mmol L^−1^ MgSO_4_, adjusted to pH 7.8; Cold Spring Harbor Protocols, 2012) enriched with macronutrients (nitrogen, phosphorus) and micronutrients (iron, trace metals, vitamins) following Jakob et al. (2018). This ASW contains no added bicarbonate or boron, so its alkalinity is low, and calcium is supplied as CaCl_2_.

### 2.2 Calcium sources and concrete characterization

Two supplementary sources of calcium were tested: reagent-grade hydrated lime (Ca(OH)_2_) and ground waste concrete. Calcium hydroxide powder (Ca(OH)₂, lab grade, purity >95%) was purchased from Lab Alley (USA). The concrete powder was produced from laboratory-prepared concrete specimens with a controlled mix design consisting of 196 kg/m³ of water, 436 kg/m³ of cement, 698 kg/m³ of fine aggregate, and 1046 kg/m³ of coarse aggregate. After three months of curing, the concrete specimens were crushed using a jaw crusher. The fine fraction of the crushed concrete was further ground using a disc mill, and then sieved through a 100 µm mesh to obtain the final concrete powder. Characterization by thermogravimetric analysis (TGA) was performed using a TA Instruments TGA 55 in a platinum crucible. The samples were heated from 40°C to 1,000 °C at a rate of 10 ° C/min under a nitrogen atmosphere with a purge flow rate of 50 mL/min. The calcium carbonate content was determined from the mass loss between 600 and 770 °C. The results for the concrete powder indicated 7.5% (w/w) portlandite (Ca(OH)_2_) and 9% (w/w) calcite. A sample of cement kiln dust was also screened but was found by TGA to be fully carbonated and was not used. Hydrated lime was added at 0.15 g L^−1^ (2.0 mmol L^−1^ Ca^2+^, 4.0 mmol L^−1^ added alkalinity). Concrete was added at 2 g L^−1^, a dose chosen to deliver the same mass of Ca(OH)_2_ (0.15 g L^−1^); this dose also introduced 0.18 g L^−1^ of calcite.

### 2.3 Experimental design

We used a 3 × 2 factorial design (Table 2) crossing calcium source (ASW alone; ASW + hydrated lime, LIM; ASW + concrete powder, CON) with the presence or absence of algae (suffix 1 or 0), with three replicate cultures per treatment (18 cultures in total). Independent variables were medium composition and the presence of algae. Response variables were pH, dissolved nitrate and calcium concentrations, and the mass of calcite at the end of the experiment; equilibrium dissolved inorganic carbon and calcite saturation state were calculated (Section 2.8). Coccolithophore strain, initial culture volume, temperature and light were held as constant as the greenhouse setting allowed.

**Table 2.** Factorial design of the experiment.

|  | **Artificial seawater (ASW)** | **ASW + hydrated lime (LIM)** | **ASW + concrete powder (CON)** |
|---|---|---|---|
| Algae absent (0) | ASW0 | LIM0 | CON0 |
| Algae present (1) | ASW1 | LIM1 | CON1 |

*Note*: Hydrated lime was added at 0.15 g L<sup>−1</sup>; concrete powder at 2 g L<sup>−1</sup> (containing 0.15 g L<sup>−1</sup> Ca(OH)<sub>2</sub> and 0.18 g L<sup>−1</sup> CaCO<sub>3</sub>). Each treatment had three replicate 1-L cultures.

### 2.4 Culture conditions and sampling

Eighteen 1-L culture bottles were filled with enriched ASW, and hydrated lime or concrete powder was added to the corresponding bottles. Bottles were shaken and left uncapped for two days to equilibrate with the atmosphere. On day 1 (13 December 2024), the LIM1, CON1 and ASW1 replicates were each inoculated with 50 mL of the same exponentially growing stock culture. Cultures were then transferred to the 6^th^ Street Rooftop Greenhouse at the University of Arizona (Tucson, AZ, USA) under natural light, and aerated with ambient air delivered by a diaphragm pump through a bubbler in each bottle. Bottles were covered with aluminum foil to limit evaporation. Air temperature in the greenhouse at the time of sampling ranged from 22.5 to 30 °C (mean ≈ 25 °C).

Cultures were sampled on days 1, 5, 8, 11, 15, 18, 21 and 26. At each sampling session, pH was measured in every bottle with a benchtop pH meter, bottles were shaken, and a 35-mL sample was withdrawn and filtered through a 0.45-µm syringe filter. Filtrates and filters were stored separately at 4 °C until analysis. On days 1 and 26, an additional 35-mL sample was frozen as an archive. On day 32, a 35-mL sample was taken from each bottle for microscopic examination. To quantify the evaporation rate, the liquid level was marked on each bottle on day 36 and again on day 41, when the experiment was terminated.

### 2.5 Chemical and mineral analyses

Filtrates were analyzed by the Arizona Laboratory for Emerging Contaminants (ALEC, University of Arizona) for dissolved Ca and K by inductively coupled plasma mass spectrometry (ICP-MS; Agilent 7700, monitoring ^44^Ca and ^39^K) after ∼100-fold dilution, and for NO_3_^−^ and PO_4_^3−^ by ion chromatography. Phosphate was frequently below the detection limit and was not used. Particulate material retained on the filters was acid-digested and analyzed for Ca and K by ICP-MS (Agilent 7700x); four unused filters were processed as blanks. The mean blank (3.7 µg Ca and 3.5 µg K per filter) was subtracted, and particulate Ca was expressed per liter of culture (35 mL filtered). Part of the Ca on each filter comes from medium retained in the filter rather than from particles; this contribution was removed by scaling the particulate K by the dissolved Ca/K ratio of the same sample, since K is not significantly incorporated into calcite. At the end of the experiment, the remaining volume of each culture was filtered, and the solids were dried and analyzed by TGA, heating to 1000 °C; the calcite content of each sample was determined from the mass loss associated with calcite decarbonation.

### 2.6 Correction of concentrations for evaporation

Because evaporation concentrates solutes, measured concentrations were corrected using the volume loss measured between days 36 and 41, which gave a final evaporation rate e_final_ for each culture (Wiegmann et al., 2018). Evaporation was assumed to increase linearly with time as culture volume decreased, so that the evaporation rate between sessions *n*−1 and *n* was

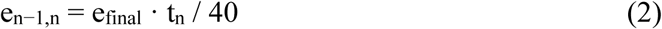

where t_n_ is the day of session *n* and 40 d is the time between the first and last volume measurements. The volume V_n_ at each session was reconstructed backwards from the volume measured on day 36 as

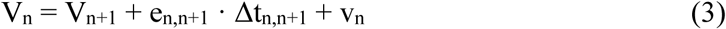

where Δt_n,n+1_ is the time between sessions and v_n_ the volume sampled at session *n* (70 mL on days 1 and 26, 35 mL otherwise, and 35 mL on day 32). The cumulative evaporation between two sessions was obtained by integrating the linear evaporation rate. The reconstructed day-1 volumes (912–1165 mL; mean 1041 mL) were consistent with the known initial volumes (1000 mL without algae, 1050 mL with algae). Sampling removes solution without changing its concentration; evaporation concentrates solutes by the factor (V_k_ − v_k_)/V_k+1_ between sessions *k* and *k*+1. A concentration X_n_ measured at session *n* was therefore corrected as

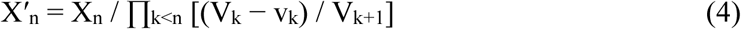

The cumulative correction was small (concentration factors of 1.05–1.12 by day 26). As an independent check, we examined dissolved K^+^, which is little affected by algal uptake at the biomass reached here. Measured K^+^ increased by 11-40% between days 1 and 26, more than the evaporation model predicts. This increase, however, is best explained by analytical drift. Indeed, day-1 readings were 15-26% below the nominal concentration of the ASW recipe (391 mg L^−1^); day-26 readings were close to the nominal concentration multiplied by the modeled evaporation factor. K^+^ was therefore not used as a tracer, and all dissolved concentrations were corrected with Equation 4.

### 2.7 Estimation of algal growth rate

Because direct cell counts proved unreliable (Section 4.4), we inferred the initial exponential growth rate of algal biomass from nitrate drawdown. Assuming that nitrate uptake is proportional to biomass *B*(*t*) = *B*_0_ e*^rt^* and to nitrate concentration *N,* with biomass-specific uptake coefficient *v*,

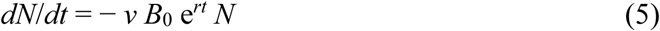

Integrating from *t* = 0 gives

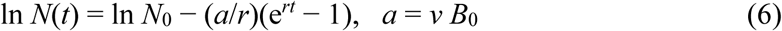

Because *v* and *B*_0_ appear only as their product, *a* is estimated as a single parameter; neither is needed to estimate *r*. Equation 6 was fitted to the evaporation-corrected nitrate data by nonlinear least squares on ln *N*, so that measurement error was treated as multiplicative. The model was fitted jointly to all nine algal cultures, with a separate initial concentration *N*_0_ for each culture and separate values of *a* and *r* for each treatment (15 parameters, 36 observations). Time was counted from inoculation. The fit was performed over the initial phase of nitrate decrease, from day 1 to 11; later data were excluded because the model cannot represent the rise of nitrate in all treatments at least between days 15 and 18 (Section 3.1). Confidence intervals (95%) for *r* were obtained by profile likelihood, using the *F* distribution. Differences in *r* among treatments were tested by comparing the full model with models in which two or all three treatments shared a common *r*, using extra-sum-of-squares *F* tests. Fits to individual cultures were used as diagnostics.

### 2.8 Carbonate chemistry and calcite saturation state

Dissolved inorganic carbon (DIC) and total alkalinity (TA) were not measured, so carbonate chemistry was calculated assuming equilibrium with the atmosphere at pCO_2_ = 420 µatm. The CO_2_ solubility coefficient was K_0_ = 0.029 mol kg^−1^ atm^−1^ (Weiss, 1974), giving [CO_2(aq)_] = K_0_ pCO_2_ = 12.2 µmol kg^−1^. Dissociation constants at 25 °C and salinity 32, the salinity of the ASW recipe (total pH scale), were taken from Lueker et al. (2000) for carbonic acid (pK_1_ = 5.859, pK_2_ = 8.997) and Millero (1995) for water, as in CO2SYS (Lewis & Wallace, 1998); the ASW contains no boron, so borate alkalinity was omitted. For a given TA, pH and DIC at equilibrium were obtained by solving

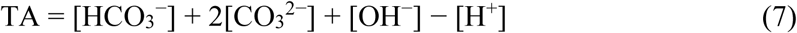

for [H^+^], with [HCO_3_^−^] = K_1_[CO_2(aq)_]/[H^+^] and [CO_3_^2−^] = K_2_[HCO_3_^−^]/[H^+^]. The TA of the LIM and CON media was taken as TA_ASW_ + 2 × 1.98 mmol kg^−1^, the alkalinity delivered by 0.15 g L^−1^ Ca(OH)_2_, assuming that concrete released the same amount of Ca(OH)_2_ as reagent lime.

TA_ASW_ was not measured. The ASW recipe contains no bicarbonate, so its nominal TA is close to zero; the mean pH of the ASW1 cultures on days 5-26 (7.18) implies TA_ASW_ = 0.26 mmol kg^−1^ at equilibrium, presumably contributed by the nutrient enrichment and the inoculum. Both values were considered. The difference in equilibrium DIC between amended and unamended media, ΔDIC_eq_, measures the additional atmospheric CO_2_ uptake capacity created by the amendment. The calcite saturation state was calculated as

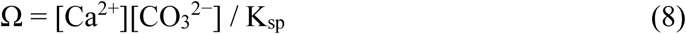

with the stoichiometric solubility product of calcite in seawater at 25 °C and salinity 35, K_sp_ = 4.6 × 10^−7^ mol^2^ kg^−2^ (Ingle et al., 1973). Ω at equilibrium at the start of the experiment was calculated from TA and the measured day-1 [Ca^2+^]. Ω at the end of the experiment was calculated for each culture from the measured day-26 pH and [Ca^2+^], with [CO_3_^2−^] from equilibrium with the atmosphere at the measured pH; this assumes near-equilibrium, which is supported by the stability of pH from day 5 onward. Ω > 1 indicates supersaturation, favoring precipitation; Ω < 1 indicates undersaturation.

### 2.9 Calcite production

At the end of the experiment (day 41), the remaining volume of each culture was measured and filtered, and the mass of calcite in the recovered solids was computed from their dry mass and TGA calcite fraction. Material adhering to the bottle walls was not recovered. For CON cultures, the 0.18 g of calcite introduced with the concrete powder was subtracted. The calcite production rate was calculated by dividing the calcite mass by the culture volume on day 41 and by the 40-day duration of the experiment, and the maximum abiotic calcite precipitation rate expected in seawater at 25 °C (0.1 mg CaCO_3_ L^−1^ d^−1^; Devriendt et al., 2021) was subtracted. Because wall growth and adhered sediment were not included, these rates are lower bounds.

### 2.10 Statistical analysis

For each treatment, means and standard deviations were computed across the three replicate cultures. Treatment means were compared with two-sample Student’s *t*-tests assuming equal variances, and changes over time within a treatment were tested by comparing day 1 with day 26. For each response variable, the three pairwise comparisons among algal treatments were adjusted for multiple testing with Holm’s method (Holm, 1979); reported *p*-values for these comparisons are adjusted. pH on days 5-26 was analyzed with a linear mixed model with treatment and sampling day as fixed effects and a random intercept for each culture, fitted by restricted maximum likelihood; treatment contrasts were tested with *t* statistics on 6 degrees of freedom (nine cultures minus three treatments). Because the design is balanced, this analysis is equivalent to a one-way ANOVA on per-culture mean pH, which was used as a check (with Tukey’s HSD). Calcite saturation states, which span two orders of magnitude, were compared on a log_10_ scale. Because the no-algae controls became contaminated after day 5 (Section 3.1), they were used only for comparisons on day 1. Analyses and graphs were produced in R (RStudio; readxl, dplyr, ggplot2) and Python 3 (NumPy, SciPy, Matplotlib).

## 3 RESULTS

### 3.1 Neither lime nor concrete inhibited nitrate uptake (P1)

Visible algal growth appeared after day 5 in some of the no-algae controls, and microscopic examination on day 32 confirmed the presence of coccolithophore cells in all “0” treatments examined. The no-algae controls were therefore excluded from analyses beyond day 1.

Initial nitrate concentrations did not differ between treatments (383-428 µmol L^−1^; all *p* > 0.6). Nitrate declined in all three algal treatments during the first 11 days, by 24% in ASW1 (*p* = 0.05), 19% in LIM1 (*p* = 0.007) and 24% in CON1 (*p* = 0.008; Figure 1A). In ASW1 the decline began immediately; in LIM1 and CON1 nitrate remained close to its initial value until day 5 and then fell (Figure S1). Nitrate then increased between days 15 and 18 in 17 of the 18 cultures, including all no-algae controls, before declining again. Over the whole experiment (day 1 vs. day 26), nitrate decreased by 28% in ASW1 (*p* = 0.03), 16% in LIM1 (*p* = 0.048) and 12% in CON1 (*p* = 0.07). By day 26, nitrate was lower in ASW1 (274 µmol L^−1^) than in CON1 (373 µmol L^−1^; *p* = 0.03) and LIM1 (360 µmol L^−1^; *p* = 0.055).

**Figure 1.**
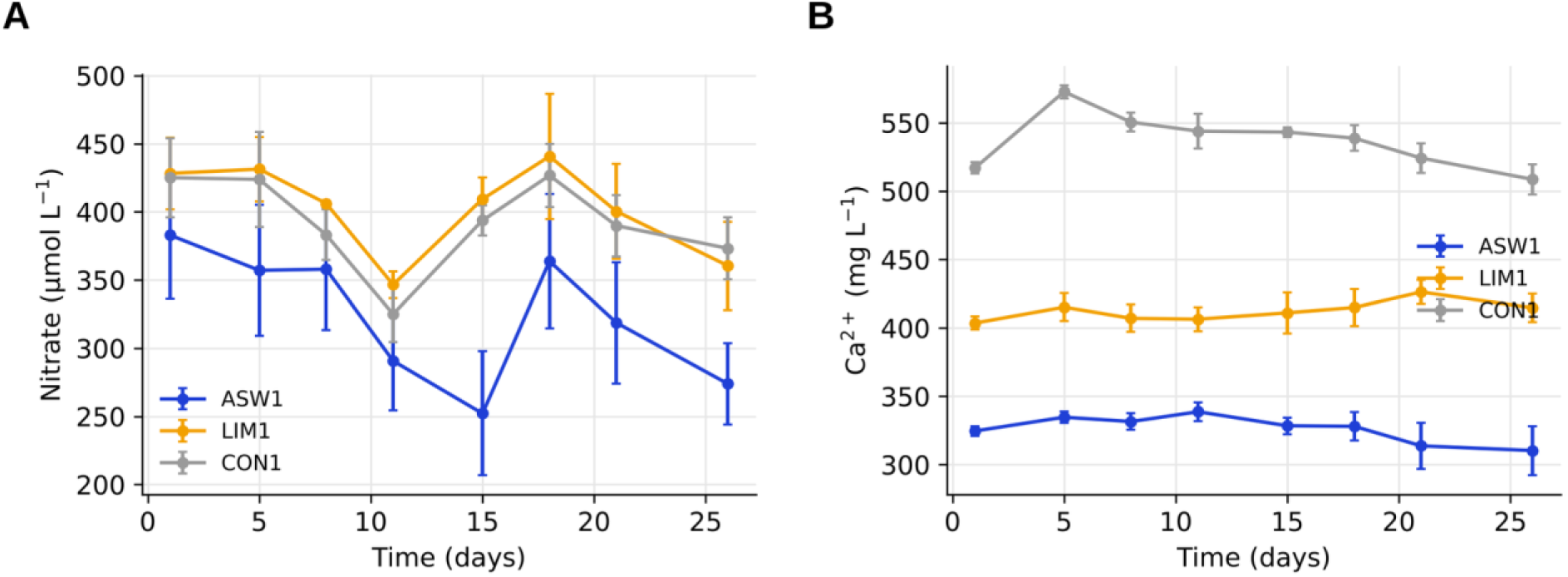
Treatment effect on nitrate and calcium dynamics. Algal cultures were grown in artificial seawater (ASW1), seawater supplemented with hydrated lime (LIM1) or with concrete powder (CON1). (A) Nitrate concentration over time, corrected for evaporation (Equation 4). (B) Dissolved calcium concentration over time, corrected for evaporation (Equation 4). Symbols and error bars show the mean ± SD of three replicate cultures.

Initial dissolved calcium was higher in LIM1 (404 mg L^−1^) and CON1 (517 mg L^−1^) than in ASW1 (325 mg L^−1^; both *p* < 0.001), and higher in CON1 than in LIM1 (*p* < 0.001). Calcium in CON1 increased further between days 1 and 5 (to 573 mg L^−1^), consistent with continued dissolution of calcium phases in concrete other than Ca(OH)_2_, including calcite (Figure 1B). Between days 1 and 26, evaporation-corrected calcium did not change significantly in any algal treatment (ASW1, −4.4 ± 5.2%; LIM1, +2.8 ± 3.8%; CON1, −1.6 ± 1.9%; all *p* ≥ 0.17), and the changes did not differ between treatments (all *p* ≥ 0.38). This is consistent with the amounts of calcite recovered at the end of the experiment (Section 3.4), which correspond to a decline of only 3-9% of the initial dissolved calcium, within the variability among replicate cultures. Calcium data for the CON0 controls after day 1 were not used.

Suspended particulate calcium, after removal of the contribution of retained medium, did not increase over the experiment (Figure S2B). It remained near zero in ASW1 (0.1-0.2 mg L^−1^) and approximately constant in LIM1 (2.4 mg L^−1^ on day 1, 3.1 mg L^−1^ on day 26), with similar values in LIM0, consistent with residual undissolved lime rather than newly formed calcite. In CON1, particulate calcium was highly variable among replicates (3.1 mg L^−1^ on day 1, 8.1 ± 5.3 mg L^−1^ on day 26).

### 3.2 Lime and concrete raised pH but did not detectably change algal growth rate (P2)

On day 1, pH was 9.4-9.6 in LIM1, 8.9-9.2 in CON1 and 7.0-7.6 in ASW1 (Figure 2A), reflecting the release of hydroxide ions by Ca(OH)_2_ in both the lime and concrete treatments; pH was higher in LIM1 than in CON1 (*p* = 0.012) and higher in both than in ASW1 (both *p* < 0.002). pH in LIM1 and CON1 fell to ≈8.1 by day 5, consistent with the invasion of atmospheric CO_2_ into the alkalinized medium, and then remained stable until day 26. Over days 5-26, the mixed model showed that pH was higher than in ASW1 (7.18) by 0.87 units in LIM1 (8.05) and by 0.93 units in CON1 (8.11; both *p* < 0.001), whereas LIM1 and CON1 did not differ (difference 0.07, *p* = 0.22). Variation among replicate cultures within a treatment was negligible compared with variation among sampling days (intraclass correlation < 0.01). An ANOVA on per-culture mean pH gave the same conclusions (*F*_2,6_ = 1025, *p* < 0.001; Tukey: LIM1 and CON1 vs. ASW1, *p* < 0.001; LIM1 vs. CON1, *p* = 0.06).

**Figure 2.**
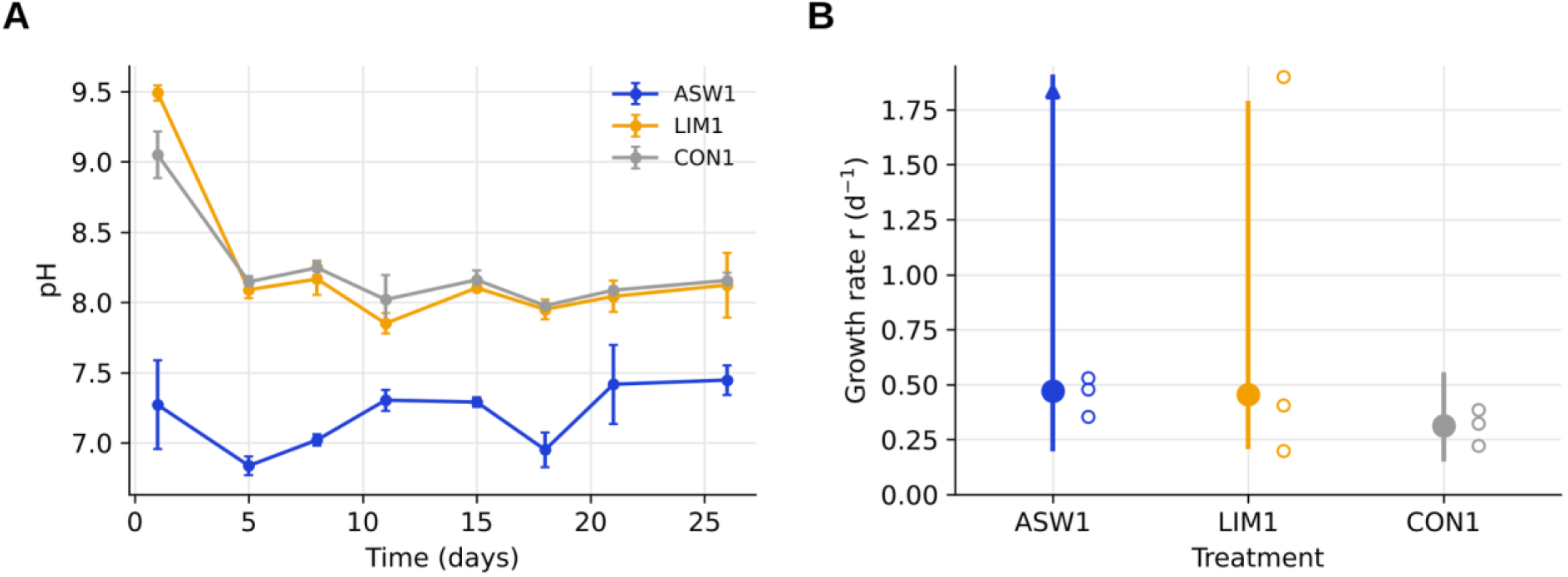
Treatment effect on pH and biomass growth. (A) pH over time in the ASW1, LIM1 and CON1 treatments (mean ± SD, *n* = 3 cultures). (B) Growth rate *r* of algal biomass inferred from nitrate drawdown over days 1-11 (Equation 6). Filled circles and vertical lines show estimates and 95% profile-likelihood confidence intervals from the joint fit to all cultures; the arrow indicates an upper limit that could not be bounded. Open circles show fits to individual cultures; the estimate for LIM1_R1 was unbounded and is plotted at the top of the axis.

The exact uptake model (Equation 6) described the nitrate data well over days 1-11 (residual SD of ln N = 0.035; Figure S1). Estimated growth rates were similar across treatments: *r* = 0.47 d^−1^ in ASW1 (95% CI 0.21 to > 4), 0.46 d^−1^ in LIM1 (0.22-1.78) and 0.31 d^−1^ in CON1 (0.16-0.55; Figure 2B). There was no evidence that *r* differed among treatments (*F*_2,21_ = 0.52, *p* = 0.60; pairwise tests, all *p* = 1.00). Confidence intervals were wide because each culture contributed only four time points to the fit; with only these points, the upper limit for ASW1 could not be bounded, and the fit for one culture (LIM1_R1) did not constrain *r* on its own. As a sensitivity analysis, normalizing nitrate to K^+^ rather than to the evaporation model (Section 2.6) gave lower but better-constrained growth rates (ASW1, 0.20 d^−1^, 95% CI 0.05-0.40; LIM1, 0.23 d^−1^, 0.09-0.45; CON1, 0.18 d^−1^, 0.06-0.33), which again did not differ among treatments (*F*_2,21_ = 0.12, *p* = 0.89).

### 3.3 Lime and concrete increased the CO_2_ uptake capacity of the medium (P3)

The amendments added 1.98 mmol kg^−1^ Ca(OH)_2_, raising TA by 3.96 mmol kg^−1^. At equilibrium with the atmosphere, this raised DIC from 0.02 to 3.45 mmol kg^−1^ if TA_ASW_ = 0, and from 0.27 to 3.66 mmol kg^−1^ if TA_ASW_ = 0.26 mmol kg^−1^ (Table 4). The additional CO_2_ uptake capacity was thus ΔDIC_eq_ = 3.4 mmol kg^−1^ in both cases, or 1.7 mol CO_2_ per mol of Ca(OH)_2_ added. The equilibrium pH predicted for the amended media (8.24-8.26) was slightly higher than that measured in LIM1 and CON1 on day 5 (8.09-8.15), consistent with the onset of calcification, which consumes alkalinity. With measured day-1 [Ca^2+^], the amended media were strongly supersaturated with respect to calcite at equilibrium (Ω = 11-16), whereas the unamended ASW was undersaturated (Ω ≤ 0.07). Because calcification consumes two moles of alkalinity per mole of CaCO_3_, the alkalinity of the unamended ASW (≤ 0.26 mmol kg^−1^) could support the formation of at most 0.13 mmol kg^−1^ of calcite, compared with up to 2.1 mmol kg^−1^ in the amended media.

### 3.4 Lime increased calcite production, and lime and concrete maintained calcite supersaturation (P4)

Calcite was recovered from all algal cultures at the end of the experiment (Figure 3A). After correction for calcite added with concrete and for abiotic precipitation, the calcite production rate was 4.5 ± 0.7 mg L^−1^ d^−1^ in LIM1, 3.6 times higher than in ASW1 (1.2 ± 0.5 mg L^−1^ d^−1^; *p* = 0.009). Rates in CON1 were highly variable (0.7, 1.4 and 17.3 mg L^−1^ d^−1^; mean 6.5, median 1.4 mg L^−1^ d^−1^) and did not differ significantly from ASW1 or LIM1 (both *p* = 0.78), likely because solids recovery was incomplete in two of the three CON1 cultures.

**Figure 3.**
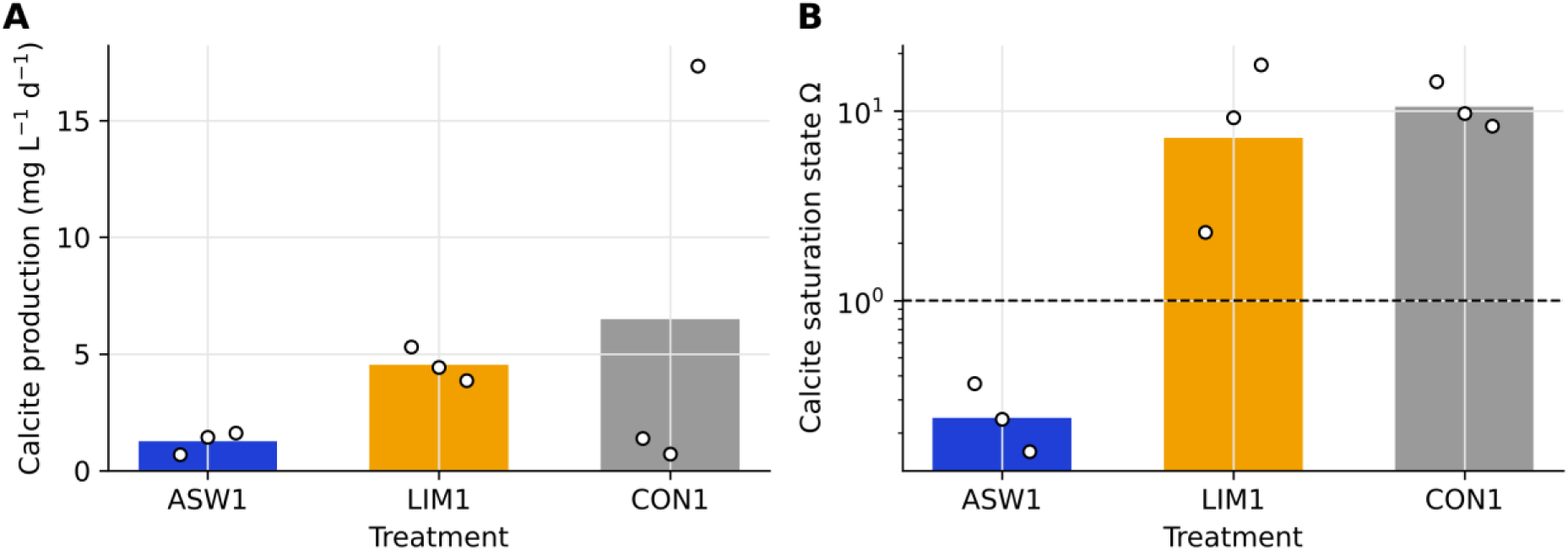
Treatment effect on clacite production and saturation state. (A) Calcite production rate in each algal culture, corrected for calcite introduced with concrete and for maximum abiotic precipitation (Section 2.9); bars show treatment means. (B) Calcite saturation state (Ω) at the end of the experiment (day 26), calculated from measured pH and [Ca^2+^] (Section 2.8); bars show geometric means (log scale); the dashed line marks Ω = 1. Open circles show individual cultures.

At the end of the experiment, ASW1 cultures were undersaturated with respect to calcite (geometric mean Ω = 0.24, range 0.16-0.36), whereas LIM1 and CON1 remained strongly supersaturated (Ω = 7.2, range 2.3-17.4, and 10.5, range 8.3-14.2, respectively; Figure 3B). Ω was higher in LIM1 (*p* = 0.012) and CON1 (*p* < 0.001) than in ASW1, and did not differ between LIM1 and CON1 (*p* = 0.57). Key results are summarized in Table 3.

**Table 3.**
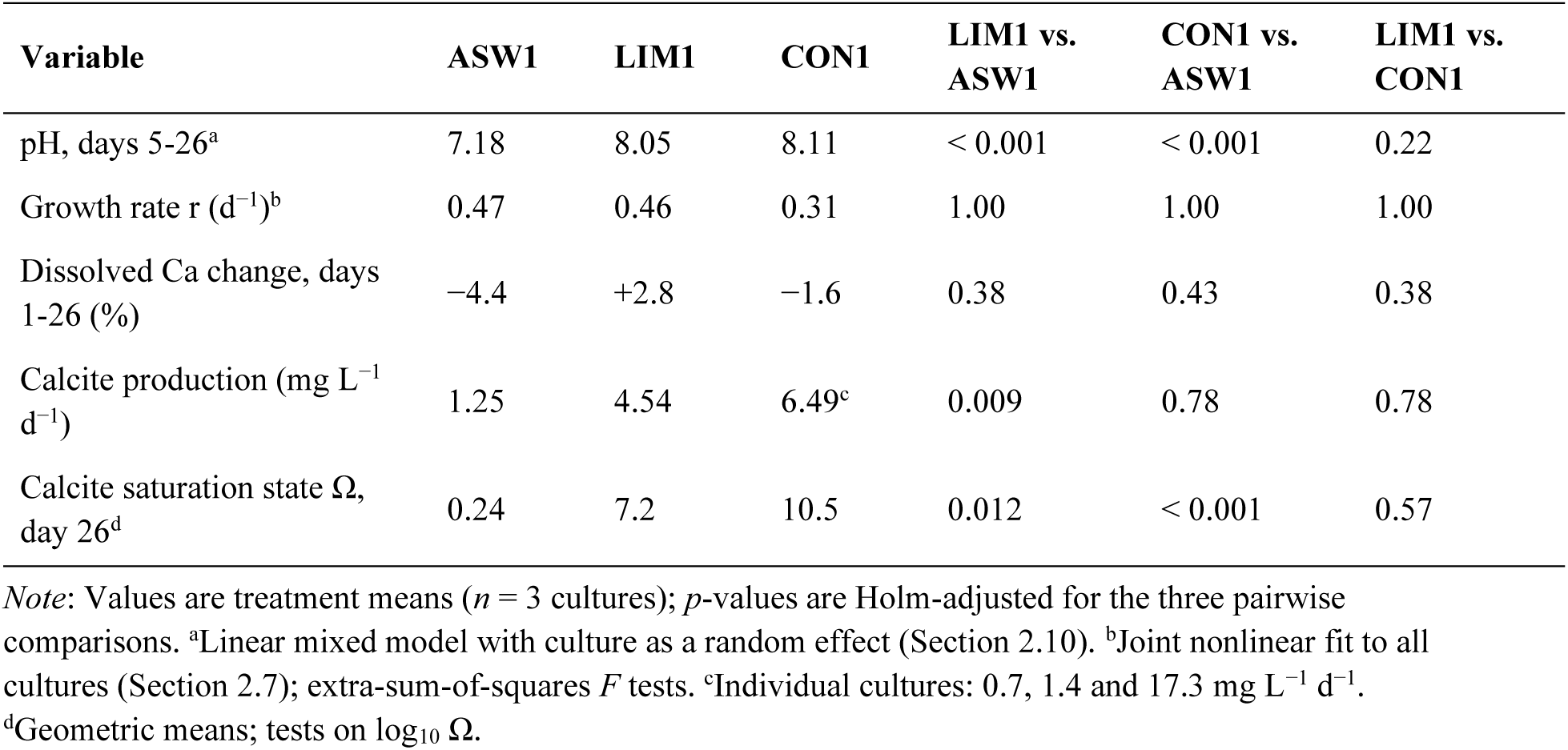
Summary of treatment effects.

| **Variable** | **ASW1** | **LIM1** | **CON1** | **LIM1 vs. ASW1** | **CON1 vs. ASW1** | **LIM1 vs. CON1** |
|---|---|---|---|---|---|---|
| pH, days 5-26<sup>a</sup> | 7.18 | 8.05 | 8.11 | < 0.001 | < 0.001 | 0.22 |
| Growth rate r (d<sup>−1</sup>)<sup>b</sup> | 0.47 | 0.46 | 0.31 | 1.00 | 1.00 | 1.00 |
| Dissolved Ca change, days 1-26 (%) | −4.4 | +2.8 | −1.6 | 0.38 | 0.43 | 0.38 |
| Calcite production (mg L<sup>−1</sup> d<sup>−1</sup>) | 1.25 | 4.54 | 6.49<sup>c</sup> | 0.009 | 0.78 | 0.78 |
| Calcite saturation state Ω, day 26<sup>d</sup> | 0.24 | 7.2 | 10.5 | 0.012 | < 0.001 | 0.57 |

*Note*: Values are treatment means (*n* = 3 cultures); *p*-values are Holm-adjusted for the three pairwise comparisons. <sup>a</sup>Linear mixed model with culture as a random effect (Section 2.10). <sup>b</sup>Joint nonlinear fit to all cultures (Section 2.7); extra-sum-of-squares *F* tests. <sup>c</sup>Individual cultures: 0.7, 1.4 and 17.3 mg L<sup>−1</sup> d<sup>−1</sup>. <sup>d</sup>Geometric means; tests on log<sub>10</sub> Ω.

**Table 4.**
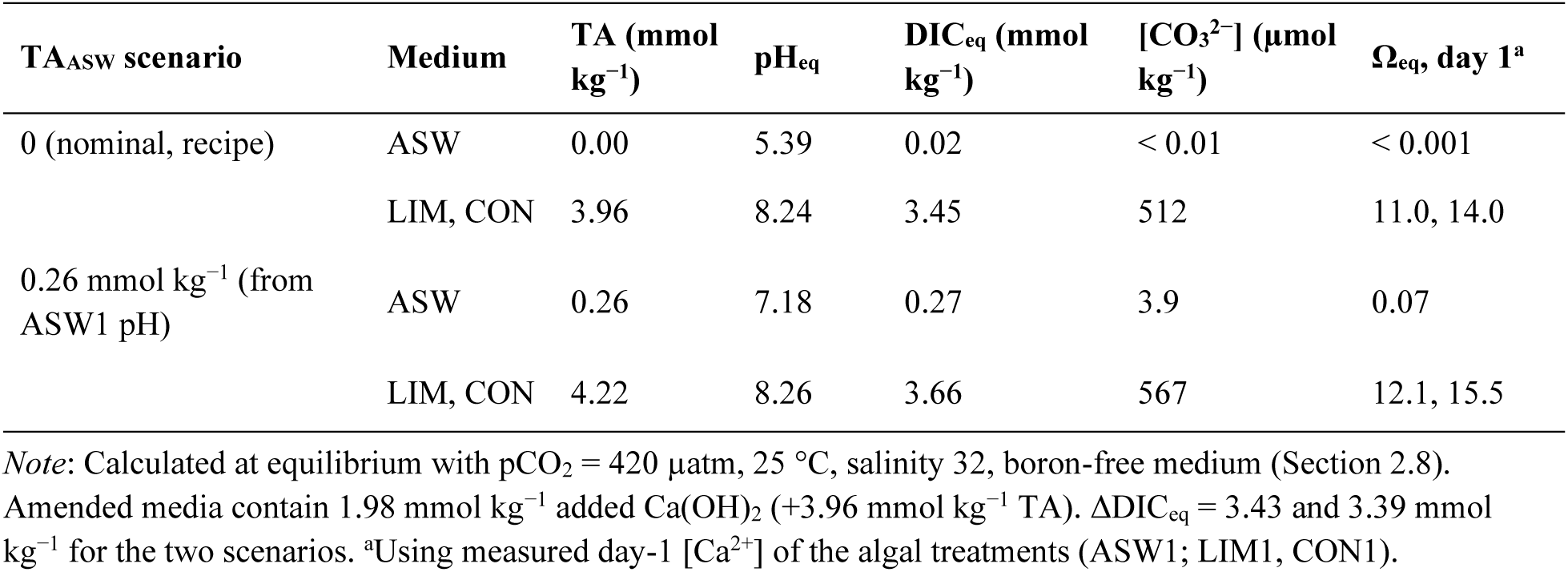
Equilibrium carbonate chemistry of unamended and amended media.

## 4 DISCUSSION

This study provides a first experimental test of using hydrated lime, alone or contained in waste concrete, as a combined source of calcium and alkalinity for coccolithophore cultivation aimed at CDR. Neither lime nor concrete inhibited nutrient uptake or calcification by *Chrysotila* sp. (P1). Both raised pH, but we found no detectable effect on growth rate, so P2 was only partly supported. Both raised the CO_2_ uptake capacity of the medium (P3) and maintained higher calcite saturation states than unamended seawater; calcite production was higher with lime, whereas results with concrete were too variable to be conclusive (P4). Concrete performed similarly to reagent lime for pH, growth and calcite saturation, indicating that the hydrated lime contained in concrete is available to the algal–carbonate system.

### 4.1 Growth and calcification with lime and concrete

The decline in nitrate, together with the accumulation of calcite, indicates that the algae grew and calcified in all treatments, in line with batch culture dynamics reported for *Emiliania huxleyi* (Perrin et al., 2016; Jakob et al., 2018). We expected higher pH to accelerate growth, because intracellular calcification produces protons that must be exported, and a higher external pH lowers the energetic cost of proton efflux. Growth rates inferred from nitrate drawdown (0.2-0.5 d^−1^, depending on the correction applied) did not differ detectably among treatments. With only four time points per culture during the initial decline, the experiment had little power to detect moderate differences in *r*. The benefit of higher pH may also accrue mainly to calcification rather than to biomass growth, consistent with the 3.6-fold higher calcite production in LIM1 (Section 3.4). The short delay before nitrate drawdown in LIM1 and CON1 (Figure S1) may reflect acclimation to the initially high pH (> 9) of these media. After the initial drop in pH, the rapid stabilization of pH in LIM1 and CON1 near 8.1 is also consistent with calcification releasing CO_2_ that is outgassed rather than accumulating and acidifying the medium.

Calcite recovered at the end of the experiment was 3.6 times higher with lime than in unamended seawater. Because calcification consumes alkalinity, the low alkalinity of the bicarbonate-free ASW limits how much calcite can form in it: at most ≈0.13 mmol kg^−1^, or ≈13 mg CaCO_3_ per culture. The 19-28 mg of calcite recovered from ASW1 cultures exceeds this limit, suggesting that part of the recovered calcite was carried over with the inoculum, which was the same in all treatments. The difference between LIM1 and ASW1 (≈3.3 mg L^−1^ d^−1^) can therefore be attributed to the alkalinity supplied by lime. In LIM1, the recovered calcite (0.84 mmol per culture on average) represents about 40% of the maximum that the added alkalinity could support (2.1 mmol kg^−1^). Recovered calcite is a lower bound, because material on the bottle walls and adhered sediment were not included in the TGA samples. Consistent with this, suspended particulate calcium did not increase during the experiment, indicating that calcite formed during the experiment did not remain in suspension but settled or accumulated on bottle walls, as commonly observed for *Chrysotila* cultures. The corresponding decline in dissolved calcium (3-9%) was not significant. These results show that in a medium lacking alkalinity, calcium supply alone (here as CaCl_2_) cannot sustain calcification; in contrast, calcification is sustained if calcium is supplied with alkalinity. Quantifying calcite in the sediment and on the walls, and measuring TA directly, will be essential in future experiments. High calcite saturation states persisted in LIM1 and CON1, in contrast to undersaturation in ASW1. This further suggests that the lime and concrete cultures could have sustained calcification for longer than the 26-day sampling period.

### 4.2 Alkalinity as the key to net carbon dioxide removal

The central problem for calcifier-based CDR is that calcification by itself releases CO_2_ (Equation 1; Frankignoulle et al., 1994). Our results point to a solution: when calcium is supplied as Ca(OH)_2_, each mole of calcium brings two moles of alkalinity, which draws two moles of CO_2_ from the atmosphere into solution. Subsequent precipitation of one mole of CaCO_3_ releases one mole of CO_2_, leaving a net removal of one mole of atmospheric CO_2_ per mole of calcite formed (Table 1). The elevated pH in the lime and concrete treatments and the equilibrium calculations are consistent with this mechanism. Indeed, the added alkalinity increases the equilibrium CO_2_ uptake capacity by ≈1.7 mol per mol of Ca(OH)_2_. The reason why this is slightly less than the theoretical 2 mol is that, at pH ≈8.25, part of the added alkalinity is carried by carbonate ions. Extrapolating the mean calcite production rate in LIM1 (4.5 mg L^−1^ d^−1^) to 1 m^3^ of culture over one year yields ≈1.66 kg of calcite, corresponding stoichiometrically to the net removal of ≈0.73 kg of CO_2_. If concrete supports the same calcite production as reagent lime, as suggested by its similar effects on pH and calcite saturation, this would require ≈29 kg of crushed concrete per cubic meter per year, renewing the 2 g L^−1^ dose every ≈25 days (14.6 batches per year). These are first-order estimates based on stoichiometry. To confirm net removal, direct measurements of CO_2_ fluxes will be required.

### 4.3 Integration with the concrete life cycle

Cement production contributes roughly 8% of global anthropogenic CO_2_ emissions, mostly from the calcination of limestone (Andrew, 2019). Coupling coccolithophore cultivation with concrete waste could contribute to decarbonizing this sector in several ways. First, concrete waste could supply calcium and alkalinity for passive CDR from air. Second, the process could be adapted for point-source capture at cement plants, using flue gas as a CO_2_ source and cement kiln dust as an additional source of calcium and alkalinity. Third, biogenic calcite could partially substitute for mined limestone as cement feedstock; because the carbon in coccoliths originates from the atmosphere, its calcination would not add fossil CO_2_. Together with CO_2_ uptake through the carbonation of cement-based materials over their life cycle (Xi et al., 2016) and the potential of biomineralized materials in construction (Beatty et al., 2022), such a cycle could contribute to net-zero cement production. A caveat is that diverting concrete waste from natural weathering forgoes the alkalinity it would eventually deliver to the ocean; full life-cycle accounting will be needed to compare these pathways.

### 4.4 Limitations and future work

Several limitations of this exploratory experiment should be noted. (i) The no-algae controls became contaminated after day 5, which prevented us from unambiguously attributing decreases in nitrate and calcium, and calcite formation, to the inoculated algae rather than to contaminating algae or abiotic processes; we corrected calcite production using a literature-based maximum abiotic precipitation rate. Separate pH probes and sampling equipment for the two sets of treatments should be used in the future. (ii) Culture volumes were measured only at the end of the experiment, so concentrations were corrected using a modeled evaporation trajectory. K^+^, which could have served as an independent tracer of evaporation, was affected by analytical drift. Volumes should be measured at each sampling, and samples should be analyzed in randomized order with check standards. (iii) Cell counts were time-consuming and imprecise because of the small size and coccolith covering of the cells, and could be obtained for only one replicate per treatment on a single date; growth rates were therefore inferred indirectly from nitrate uptake, from only four time points per culture. A nitrate rise common to all cultures between days 15 and 18, which was not mirrored by K^+^, suggests an analytical artifact in the nitrate measurements; more frequent sampling and randomized analysis order would strengthen growth-rate estimates. (iv) The high salinity of the medium hindered the analysis of phosphate. (v) DIC and Ω were calculated assuming equilibrium with the atmosphere rather than measured, and the alkalinity of the unamended medium was inferred from the recipe and measured pH; TA and DIC should be measured directly in future experiments. (vi) Material on the bottle walls and adhered sediment were not included in the TGA samples, so calcite production is underestimated; solids recovery was also difficult in two CON1 replicates. (vii) Calcite carried over with the inoculum was not quantified, and the 26-day sampling period was too short to reveal the limits of growth in ASW, where alkalinity and Ω were lowest.

Future work should optimize nutrient supply and the dose and delivery schedule of lime or concrete; Jakob et al. (2018) reported substantially higher coccolith productivities under optimized photobioreactor conditions. Experiments should then be scaled to larger photobioreactors (> 50 L) with automated monitoring of cell density and carbonate chemistry (Apai et al., 2024). Ultimately, quantifying the fraction of atmospheric CO_2_ sequestered in biogenic calcite will require isotopic tracer experiments in controlled growth chambers.

## 5 CONCLUSIONS

Supplying calcium to coccolithophore cultures as hydrated lime, either as a reagent or contained in ground concrete waste, did not inhibit growth or calcification, raised pH and the capacity of the medium to absorb atmospheric CO_2_, and increased calcite saturation state and, with reagent lime, calcite production relative to unamended seawater. By supplying alkalinity together with calcium, this approach can offset the CO_2_ released by biocalcification, a key obstacle to calcifier-based CDR. These results provide proof of concept for coupling coccolithophore cultivation with the reuse of concrete waste, a pathway that could contribute both to atmospheric CO_2_ removal and to a more circular, lower-carbon cement industry.

## ACKNOWLEDGMENTS

We thank the Arizona Laboratory for Emerging Contaminants (ALEC) for chemical analyses, especially Ashlyn Zilch for all her work on our samples. M.F. thanks Regis Ferriere and Margaret Evans for support and comments on an earlier version of the manuscript. D.A., J.C. and H.J.K. lead the ATMOSPHERICA team.

## AUTHOR CONTRIBUTIONS

**Maxime Ferriere:** conceptualization, investigation, formal analysis, visualization, resources, writing – original draft. **Antonin Affholder:** conceptualization, methodology, formal analysis, writing – review and editing. **Jack Welchert:** methodology, supervision, writing – review and editing. **Matthew Katterman:** methodology, resources, writing – review and editing. **Hang Zeng:** investigation (concrete and TGA analyses), methodology, writing – review and editing. **Hee-Jeong Kim:** resources, writing – review and editing. **Daniel Apai:** conceptualization, writing – review and editing. **Joel Cuello:** resources, supervision, writing – review and editing.

## DATA AVAILABILITY STATEMENT

The data and analysis scripts (R and Python) that support the findings of this study are openly available in *add repository* at *add DOI*.

## FIGURE LEGENDS AND FIGURES

**Figure S1.**
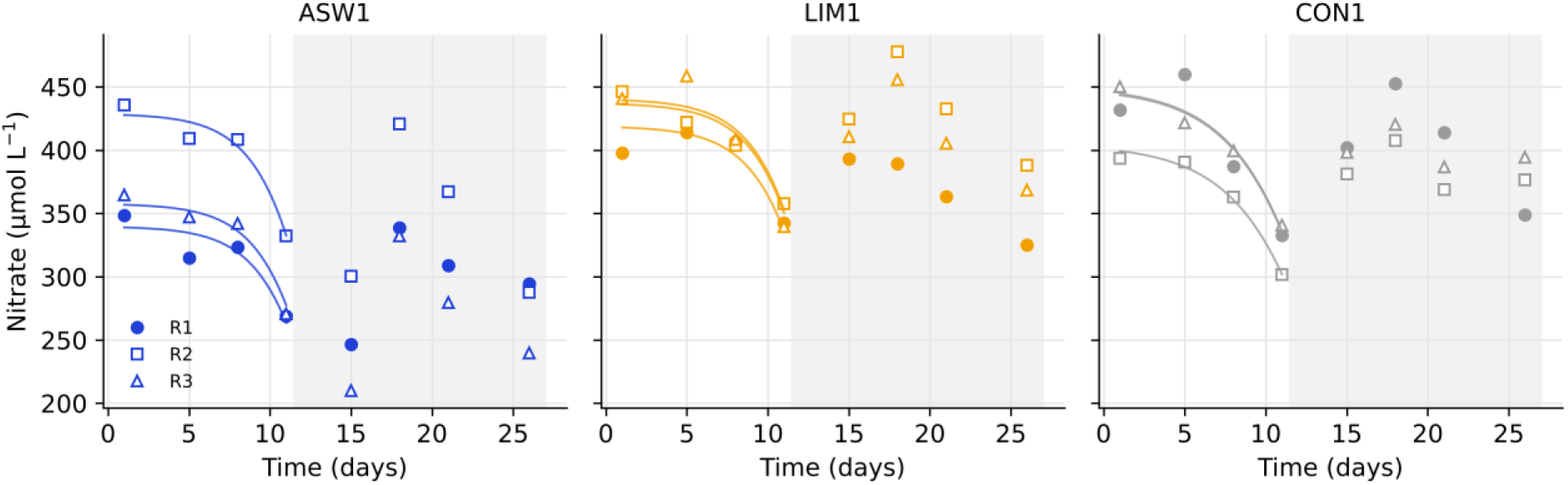
Evaporation-corrected nitrate concentration in each algal culture. Symbols indicate R1-R3 culture replicates. Lines indicate the joint fit of Equation 6 over days 1-11. Data in the shaded period (days 15-26) were not used in the fit.

**Figure S2.**
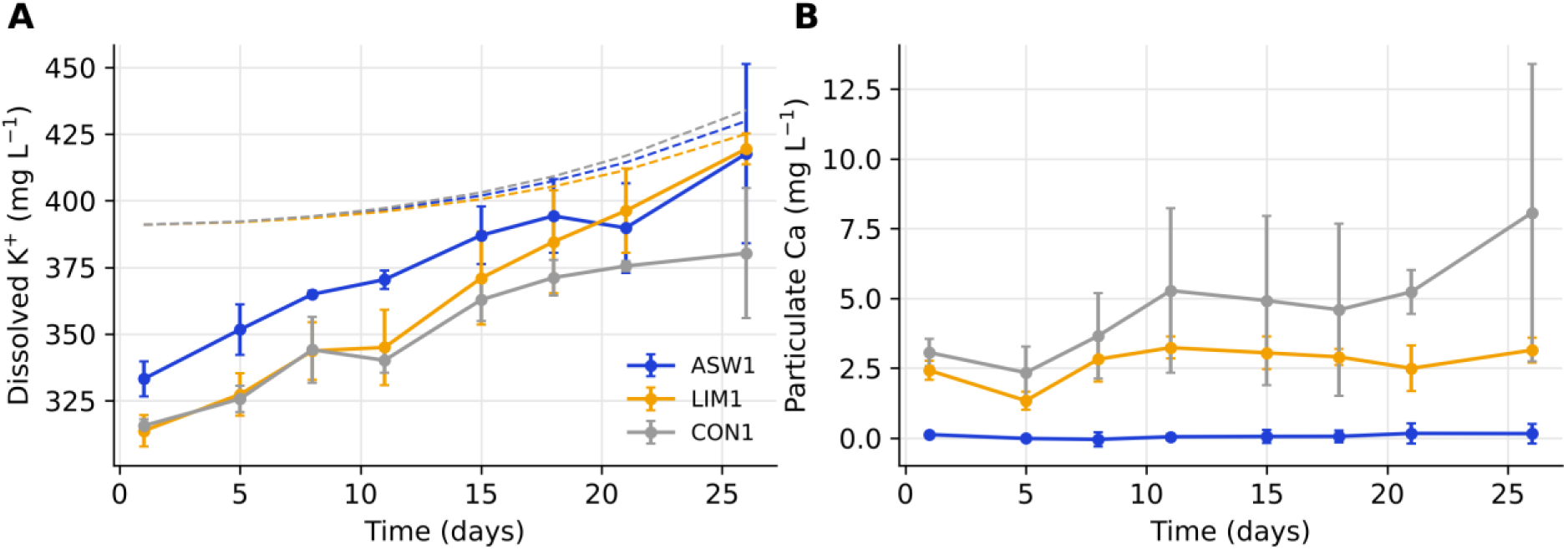
Treatment effect on dissolved potassium ions and particulate calcium. (A) Dissolved K^+^ concentration over time, uncorrected; dashed lines show the nominal concentration of the ASW recipe (391 mg L^−1^) multiplied by the modeled evaporation factor (Equation 4) for each treatment. (B) Suspended particulate calcium retained on 0.45-µm filters, after subtraction of blank filters and of calcium in retained medium (Section 2.5), per liter of culture. Symbols and error bars show the mean ± SD of three replicate cultures.

